# EEG-Based Decoding of Color and Visual Category Representations Is Reliable Within and Across Sessions

**DOI:** 10.64898/2026.01.18.699677

**Authors:** Chen Frenkel, Leon Y. Deouell

## Abstract

The human visual system represents stimuli in a rich and detailed manner. Traditional methods of studying visual representations in humans, such as event-related potentials (ERP), revealed numerous distinctions between the brain activity elicited by different categories of stimuli. However, these methods miss the information embedded in the spatial distributions of brain activity, or patterns, and are not always sensitive to study visual representations of different stimuli at the single participant or single trial level. Time-resolved multivariate pattern classification analysis (MVPA), or Decoding, efficiently extracts the visual representations of stimuli from the EEG topography without a-priori assumptions about the location of the effect in time and space at the single participant level. The rich information this method provides has increased its popularity dramatically in recent years. Yet, different participants show variable quality of decoding performance, and it is unclear if the accuracy of decoding is maintained within participants across multiple sessions, tasks, attentional conditions and visual features. In the current study, participants performed three visual tasks, over two sessions (1-7 days apart). We examined the correlation of decoding accuracy: within the cross-validation set, between sessions, between features (color and category) and to different measurements of the ERP signal and behavioral performance. We also examined how models generalized to different tasks and different attention conditions. We found that decoding accuracies varied substantially across participants, and that decoding accuracy was reliable within participant, over sessions, attention condition and task. This suggests the decodability behaves like an individual trait. Moreover, the spatial patterns underlying the decoding (classification weights) generalized across different tasks, attentional conditions and sessions. This suggests minimal representational drift at the resolution allowed by the EEG. We conclude that EEG decoding is a reliable method, and that visual representations are stable.

## Introduction

The human visual system encodes stimuli with remarkable richness and detail. Understanding the neural underpinnings of our complex visual experiences is an ongoing challenge to neuroscience. Electrophysiological methods, such as electroencephalography (EEG), offer high temporal resolution and the ability to capture dynamic perceptual processes. However, extracting the specific content of percepts from EEG signals presents significant hurdles. Traditional approaches, such as event-related potentials (ERP), often lack the sensitivity, precision and statistical power required to distinguish subtle, transient patterns that differentiate between responses to multiple visual categories (Hebart & Baker, 2018). A major impediment in this regard is the individual variability between the participant’s neural responses, both in the spatial distribution of the responses over the scalp and in its temporal dynamics (Rousselet & Pernet, 2011). That is, while the gross spatio-temporal distribution of ERP responses is similar enough across individual to allow group statistics, more subtle differences between responses to different stimuli, may be masked by averaging.

Time-resolved multivariate pattern analysis (MVPA), or decoding, has emerged as a powerful alternative for uncovering distinctions between neural responses to visual events without prior assumptions about the scalp locations or the timing of the relevant effects. By enabling individualized decoders, MVPA accommodates variability in sensor placement and individual brain anatomy, yielding robust estimates of neural distinctions. This methodological flexibility, combined with increased accessibility of MVPA techniques (e.g., Bode et al., 2019; Fahrenfort et al., 2018; Hanke et al., 2009; Meyers, 2013; Haxby et al., 2014; Oosterhof et al., 2016; Treder, 2020), has fueled a surge of interest in recent years.

With this growing popularity, substantial efforts have been devoted to optimizing MVPA analysis, including comparisons of algorithms (e.g., Grootswagers et. Al., 2017) and processing pipelines (e.g., van Driel et. Al., 2021). These studies provide insight into variability across experiments and analysis protocols. A frequently overlooked observation is that participants exhibit considerable differences not only in the patterns of brain activity, reflected in classifiers weights, but also in decoding accuracy, that is, the extent to which a machine learning algorithm can dissociate the responses to distinct categories. It is not clear whether this variability arises from session-specific factors, like measurement noise or fluctuations in alertness, or from stable individual traits, such as how strongly relevant brain regions project to EEG sensors or how vividly participants form visual representations.

To address these questions, we investigated whether decoding accuracy is consistent within participants between recording sessions separated by days. Across the two sessions, we inspect accuracy, or how well the decoding model captured the representations of each participant in each session, and the generalization between the two sessions, or how similar are the spatial representations captured by these models between sessions. We hypothesized that stable accuracy rates within and between sessions would validate decoding as a reliable method. Independently, high generalizability of classifiers across sessions would suggest that information captured by the decoding model reflects stable neural properties. The latter notion of stability, or lack thereof, is tied to the concept of “representational drift,” which describes changes in the tuning properties of single neurons with time, despite consistent behavioral outputs (Driscoll et. Al., 2022). In this context, a situation where accuracy correlation between sessions is high, but classifiers do not generalize from one session to the other would reflect a representational drift. Drift has been observed on timescales ranging from seconds to days, but recent findings using fMRI suggest that analysis of whole-brain activity may be relatively robust to such drift over short timescales (days to weeks) while being more susceptible over longer durations (Roth & Merriam, 2023). We hypothesized that MVPA applied to EEG would similarly show minor drift and high generalizability over days, reflecting stable perceptual representations in these time scales.

Much of the research into representational drift has focused on neuronal populations in animals engaged in a single task and a single visual property (Rule et. Al., 2019; Rule et. Al., 2020). Therefore, it is not clear whether different levels of the perceptual hierarchy are influenced in the same way; representational drift may differentially affect low-level (e.g., color) and high-level features (e.g., category). In the current study, we compared the generalizability at both processing levels. Additionally, it is unclear how factors such as attention or task demands interact with representations. If the task and attentional demands play a role in modulating representations, cross-task generalizability might be reduced, with decoding performance varying based on task-specific contexts in spite of consistent visual input. To investigate these possibilities, we manipulated task and feature-based attention, assessing the reliability of MVPA across both visual features and conditions to understand the interplay between representational drift, attention and task context.

## Methods

### Participants

32 participants took part in the study (16 females, aged 18-45) each participant completed two experimental sessions, separated by 1-7 days (on average 4 days, SD = 2.7). All participants reported no history of psychiatric or neurological illness and declared not taking medications affecting the nervous system at the time of the experiment. Participants were compensated with either course credit or a fixed hourly rate (the equivalent of ~$15/h). The ethics committee of the Hebrew University of Jerusalem approved the study, and all participants provided written informed consent and a screening questionnaire prior to participation. Two participants were disqualified for either not completing the full experiment or due to excessive recording noise; all further analyses are based on 30 participants (mean age = 23.9, SD = 3.5).

### Stimuli and Task

The stimuli were presented using Psychopy software (V2022; Peirce, 2007). Stimuli consisted of schematic drawings from four distinct categories (people, animals, cars, and inanimate objects; Figure 1) presented in four colors (black, white, red, and blue). Each category included 16 unique exemplars. Images were presented sequentially with predetermined statistical regularities of the categories or colors. In each experimental session one feature (color or category; *Attended Feature*) was task relevant and predictably followed a first-order sequential probability of 85%, while the other feature was presented randomly with 25% sequential probability (*Unattended feature*). Each session included four different tasks, each divided to four equal blocks separated by a short break. In each task, participants monitored the stimulus stream and responded to unexpected probe trials (every 2-15 trials, 15% of total trials). During these probed trials, the sequence was temporarily halted, and participants answered a two-alternative forced choice question about the stimulus attended feature (i.e. color or category) using the left or right arrow keys. The incorrect choice (foil) and the key mapping (i.e., which key indicated the correct answer) were randomly selected for each probed trial. This task design was chosen to maintain attention while minimizing motor preparation effects on neural representations.

**Figure 1.**
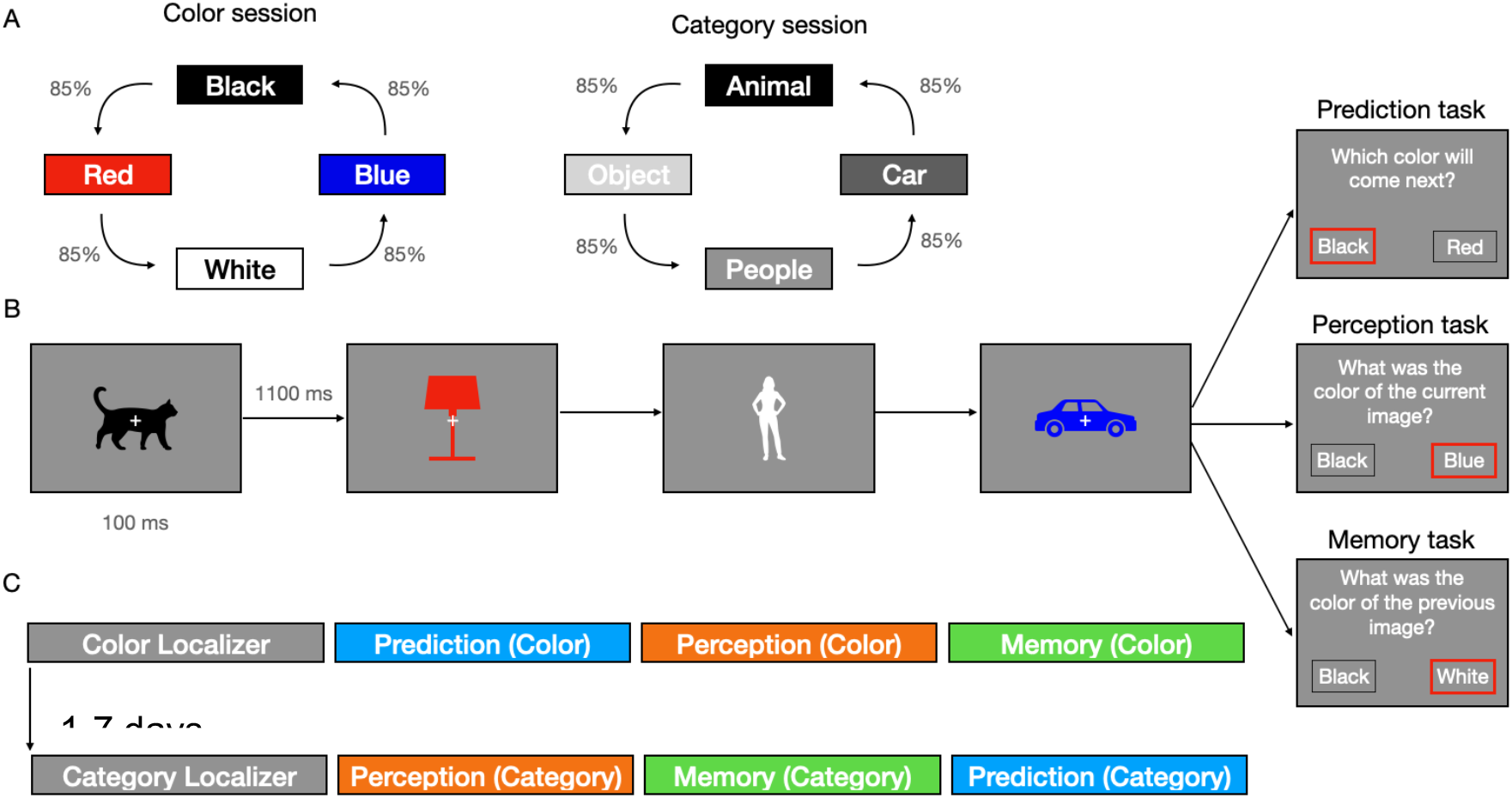
Schematic description of the task. A) An example of the sequences presented during the tasks. In each session all possible four-item circular sequences of that form were presented. One was used for practice, four were presented during the localizer task and the remaining one during the main tasks. B) The task timeline, each stimulus was presented for 100 ms, followed by a blank screen with a fixation cross for 1100 ms. Stimuli followed the sequence and every 2-15 trials (unexpectedly) a question appeared, according to the task. C) a general outline of the order. Localizer blocks always came first, followed by the main tasks in counterbalanced order (between participants).

Each session began with a baseline prediction task intended to be used for classifier training. Probe trials in this task instructed participants to predict the next trial’s color or category, based on the sequence. Each of the four blocks of this task presented a different predefined sequence of 275 images with conditional probability of 100% (i.e. with no irregular images). The sequence was presented to the participant before each block (e.g. red, white, blue, black; Figure 1). Only 10% of the images required response and were not used for training. Across the experiment every category was preceded equally by every other category, and every color was preceded by every other color. This allowed us to train classifiers for the different categories or colors minimizing confounding effect of the previous stimulus. Following the baseline task participants proceeded to perform tasks designed to manipulate the participants intention to form expectations about upcoming stimuli. In these tasks, participants were required to either predict the next item, as in the baseline task (Prediction Task), describe the current item (Perception Task) or report the previous stimulus (Memory Task). Each block of trials included probes of only one task. The rational for this manipulation pertains to questions which will be described in a separate manuscript. Each task included 4 blocks of the same sequence, which was different from the four sequences used in the baseline task. As mentioned above, each sequence included 15% violations of the sequence. Each task started with a practice block using yet another sequence, thus exhausting the 6 possible sequences of 4 options presented cyclically. Session and task orders were counter-balanced between participants to avoid any order related biases.

### EEG Recoding and Preprocessing

EEG data were recorded using a 64-channel Active II system (Biosemi, Netherlands) at a sampling rate of 1024 Hz with an online low pass filter at 1/5^th^ the sampling rate. Electrodes were placed in an elastic cap according to the extended 10-20 system. External electrodes were placed on the mastoids, nose, at the outer canthi of the eyes and below and above the left eye to monitor eye movements. EEG analysis was conducted offline using python MNE (V2022.2; Gramfort et. Al, 2014). Data was high-pass filtered with a cutoff of 0.5Hz (using a FIR non-causal filter) to eliminate low frequency trends throughout the session. The data was inspected for malfunctioning channels, and if such were found they were spherically interpolated. Data was then re-referenced to the average of all 64 channels. Finally, data was low-pass filtered at 100 Hz and a notch filter was applied around 50 Hz to remove electrical noise. Raw data was segmented (−200 to 500 ms relative to stimulus onset) and segments including an absolute minimum to maximum difference of more than 200uV were rejected from all further analyses.

We performed an ERP analysis to investigate the relationship between amplitude and variability of the ERP, and classifier accuracy. For this analysis data was additionally low-pass filtered at 40Hz and eye blink effects on the EEG were removed using ICA. Epochs were baseline corrected to 200-50 ms prior to stimulus onset and epochs with a min-max amplitude difference exceeding 200uV were removed. We used these epochs to detect the peaks of the early visual components P1, defined as the maximum amplitude in the window of 70-150 ms post stimulus onset, and N1, defined as the minimum amplitude in the window of 120-200 ms post stimulus onset. The P1 peak was always chosen so it precedes the N1 peak. Peaks were detected for every color and category separately in two visual channels – PO8 and PO7. The signal estimate was than calculated as the amplitude difference between the peaks (P1-N1), which was then averaged across channels, categories and colors to create one estimate per participant per session. Using these segments we also calculated an ERP noise estimate, by calculating the standard deviation (SD) between trials at each time point between 100-300 ms post stimulus onset and averaging these variability measures across time for each color, category and channel. The SDs were than averaged across conditions and channels and normalized to provide a single estimate per participant per session.

### Decoding Analysis

In order to model visual representations of color and category, we trained two independent multi-class classifiers, one for color and one for category for each session (altogether four classifiers) using the baseline blocks for training. Decoding was done using linear discriminant analysis (LDA, implemented using Scikit learn; Pedregosa et. Al., 2011) trained to separate stimulus category or color from the vector of amplitudes at the 64 EEG channels down sampled to 128 Hz. Decoding was done for each time point independently using a sliding estimator analysis, implemented in MNE, across the full segment length. Temporal generalization matrices (TGM; King & Dehaene, 2014) were created by applying the model of each time point to all other timepoints, also implemented in MNE. The following analyses were performed:

1. To provide an estimate of the decoding performance of each decoding model, a 6-fold cross-validation (CV) procedure was performed over the baseline data. The number of trials per label, as well as of the previous label were balanced within each CV fold to avoid biasing of the classifier.
2. To quantify generalization of the models between sessions and tasks we used the full baseline data as the training set and tested on independent data sets.
  a. To generalize between sessions the trained model of one session was applied to the other.
  b. In order to generalize across tasks, the trained model was applied to each task within the same session.

Classifier performance was assessed using the area under the receiver operator curve (AUC) using a one-vs-rest (OVR) multiclass protocol (associated with a chance level of 50%). To control for potential confounds, we excluded all irregular and question trials and trials following such an event from the decoding analysis, as these might be confounded by the response to an unpredictable event.

### Statistical Analysis

Each decoding time course and temporal generalization matrix were compared to the chance level of 0.5 using a one sample cluster permutation test with 10,000 permutations (Allefeld et. Al., 2016). Critical p value was corrected for multiple comparisons using Bonferroni correction (critical p=0.004 for 12 comparisons). ^1^

In order to assess the correlation between and within sessions across participants, the average AUC for the peak perceptual time (100-300 ms post stimulus onset) was used as the dependent variable, thus reducing each participant’s accuracy estimate to one value per condition. Pearson correlations were calculated, and their significance was tested, corrected for multiple comparisons using Bonferroni correction (critical p=0.008 for 6 correlations between decoding accuracies and 0.0125 for 4 correlations to ERP and behavior).

## Results

We recorded EEG data from 30 participants across two sessions (1-7 days apart). Participants mainly viewed and occasionally responded to visual stimuli varying in color and category. Using multivariate pattern analysis (MVPA), we trained classifiers to decode the perceived color and category of the stimuli from EEG signals. We probed test-retest reliability of individual differences in decoding accuracy, as well as possible factors contributing to individual differences. We examined decoding results and how well they generalized between sessions, tasks and attentional conditions in order to understand how stable representations are.

### Category and color classification

Four decoding models were created for each participant (attended color, attended category, unattended color, unattended category). The classifiers were trained and tested on the baseline blocks of each session, in which the participants viewed sequences of colored (red, blue, black and white) silhouettes of people, animals, objects and cars. First, we estimated each model’s ability to discriminate between the different categories and colors when these dimensions were attended. Model accuracy was based on the average of 6-fold cross validation AUC. A cluster of above chance decoding (0.5) results was detected for color from 78 - 500 ms, (p < 0.001, figure 2A) and for category from 54 - 500 ms sec (p<0.001, figure 2B).

**Figure 2.**
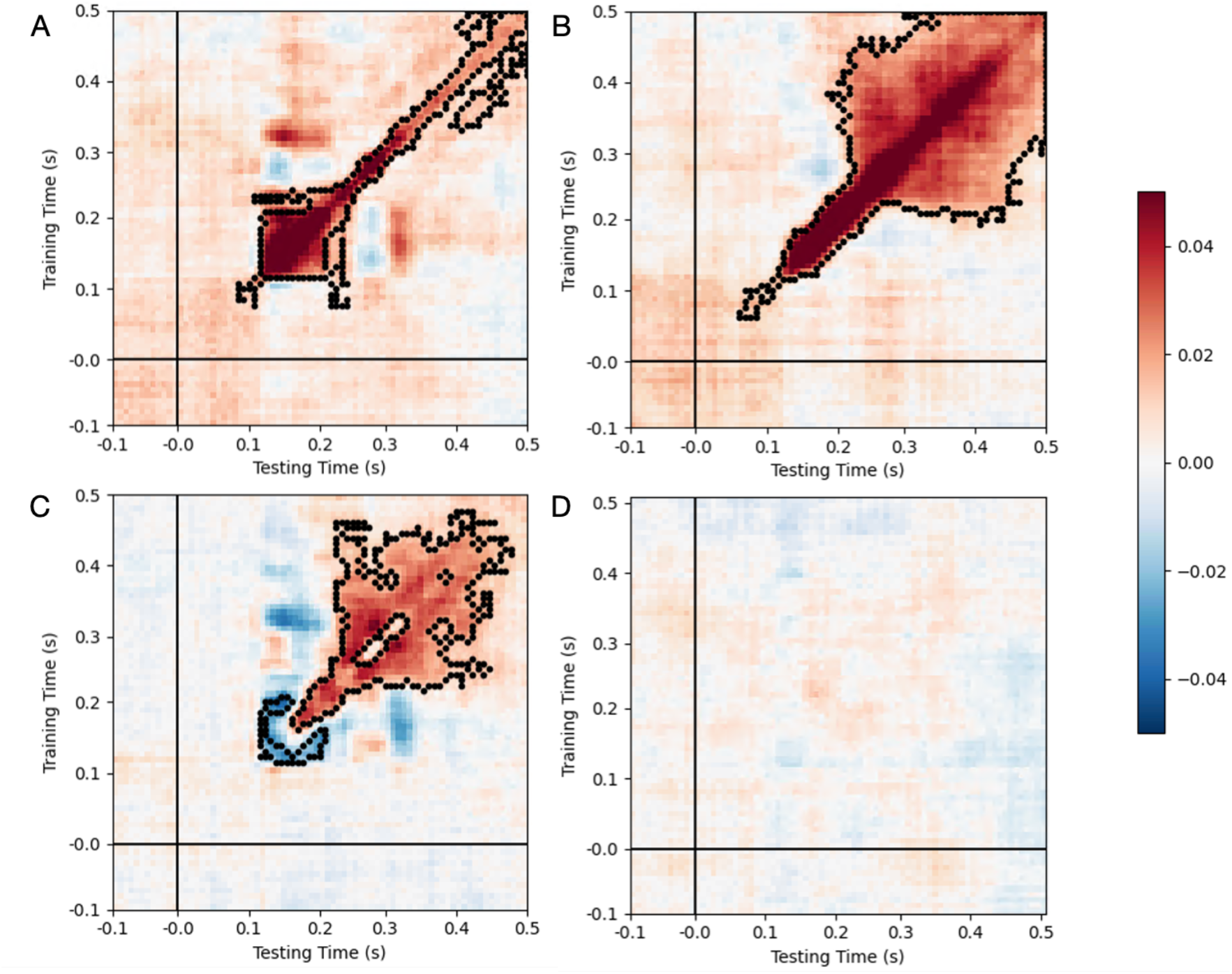
Temporal generalization matrices of the cross-validation decoding AUC averaged across participants and cross validation folds. The x axis represents the time point of the testing set and the y axis represents the time point of the training sets within the cross validation. Color reflects the AUC. A) Decoding of color identity from the predictable session. B) Decoding of category identity from the predictable session. C) The difference between the averaged color decoding in both sessions and category decoding in both sessions. D) The difference between the averaged decoding in both attended sessions and unattended sessions.

Color and category clusters appear to have a distinct pattern of temporal generalization, where color decoding generalizes early and briefly, whereas category appears to generalize after about 250ms as would be expected during these different levels of processing (Figure 2). In order to estimate the significance of these differences between color and category decoding timing we conducted a cluster permutation test on the difference between category and color TGM. Consistent with the above observation, we detected a significant early cluster where color was better decoded than category (109-203 ms p=0.003) and a later cluster where category was better decoded than color from 156 to 468 ms post stimulus onset (p<0.001). These two clusters appear to partially overlap, yet the overlapping window is associated with the end of the temporal generalization of the color model that co-occurs with the category model achieving higher accuracy along the diagonal (see Figure 2C). Finally, the results of the unattended model were qualitatively similar, and no significant differences were detected between the TGMs of the two conditions (Figure 2D). This suggests that feature-based attention did not influence the model’s ability to decode color or category of the images from EEG within this time window, possibly because the images were always task-relevant.

### Reliability of decoding accuracy

Participants varied in the accuracy of decoding (Figure 3). To understand how consistent the performance of the MVPA model is at the single participant level, we calculated an average AUC per participant in the early perceptual window (100-300 ms). We correlated the AUCs across participants on three levels: within session, between odd and even CV folds of the CV set from the attended session of each feature, between the attended and unattended sessions of each feature, and between the features, color and category, separately when attended and unattended. In all cases, significant correlation will indicate that decoding accuracy is a stable trait at the participant level. The correlation within the CV set serves as a baseline estimate, since it reflects the reliability of the CV set itself. Indeed, high and significant correlation was detected for both color (r=0.82, df=28, p<0.001) and category (r=0.75, df=28, p<0.001; Figure 3A). Despite the different attentional conditions between the two sessions, a significant correlation was also detected between the sessions for both color (r=0.54, df=28, p=0.002) and category (r=0.72, df=28, p<0.001; Figure 3B). Finally, the correlation between the decoding of color and category reflects whether decoding levels are feature specific or a more general property of the participant. A significant correlation was detected in the attended session (r=0.68, df=28, p<0.001; Figure 3C). In the unattended sessions, a marginally significant correlation was detected, not surviving the correction for multiple comparisons (r=0.46, df=28, p=0.01). These results suggest that decoding performance is stable across features, more prominently when attention is allocated to the feature.

**Figure 3.**
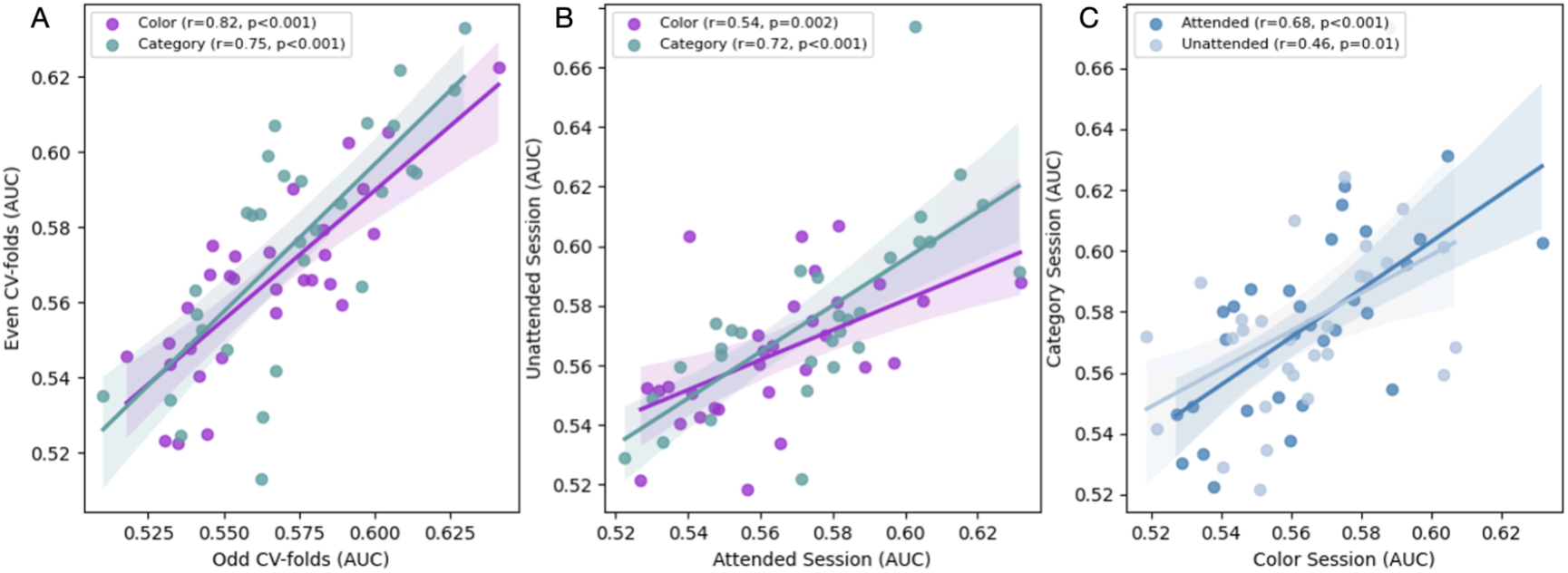
Scatter plot with linear trend of the averaged AUC of the perceptual time (0.1-0.3 seconds post stimulus onset) per participant. A) presents the linear trend within the cross validation between odd (X axis) and even (Y axis) CV folds separately for color and category (marked by color). B) presents the linear trend between the attended session (X axis) and the unattended session (Y axis) separately for color and category. C) Presents the linear trend between decoding of color (X axis) and category (Y axis) for the attended and unattended session (marked by shade).

### Classifier generalization across sessions and tasks

The above results show that single participant decodability is a relatively stable trait of participants – a participant who yields relatively high decoding accuracy is likely to show relatively high decoding accuracy in different sessions and for different features. However, this reliability of decoding accuracy was shown when classifiers were trained and tested using cross validation within each session or task. Therefore, it does not necessarily entail the stability of the underlying neural representations, captured by the classifiers. That is, one might represent a category in a different way on different days, both equally well decodable. To address the question of representational stability, we used the trained classifier of the attended session and tested it on data from the unattended session for both color and category identity. Although the classifiers were trained in one session and tested on EEG data from another session, collected on a different day, with reapplication of the electrodes, a significant cluster was detected for both color (Range: 86 - 359 ms, p<0.001; Figure 4A) and category (Range: 109 - 484 ms, p<0.001; Figure 4B). The clusters were consistent in the temporal generalization patterns with the ones detected in the CV within session (shown in Figure 2), suggesting that there is a robust and consistent activity pattern that can be transferred between sessions in the range of days.

**Figure 4.**
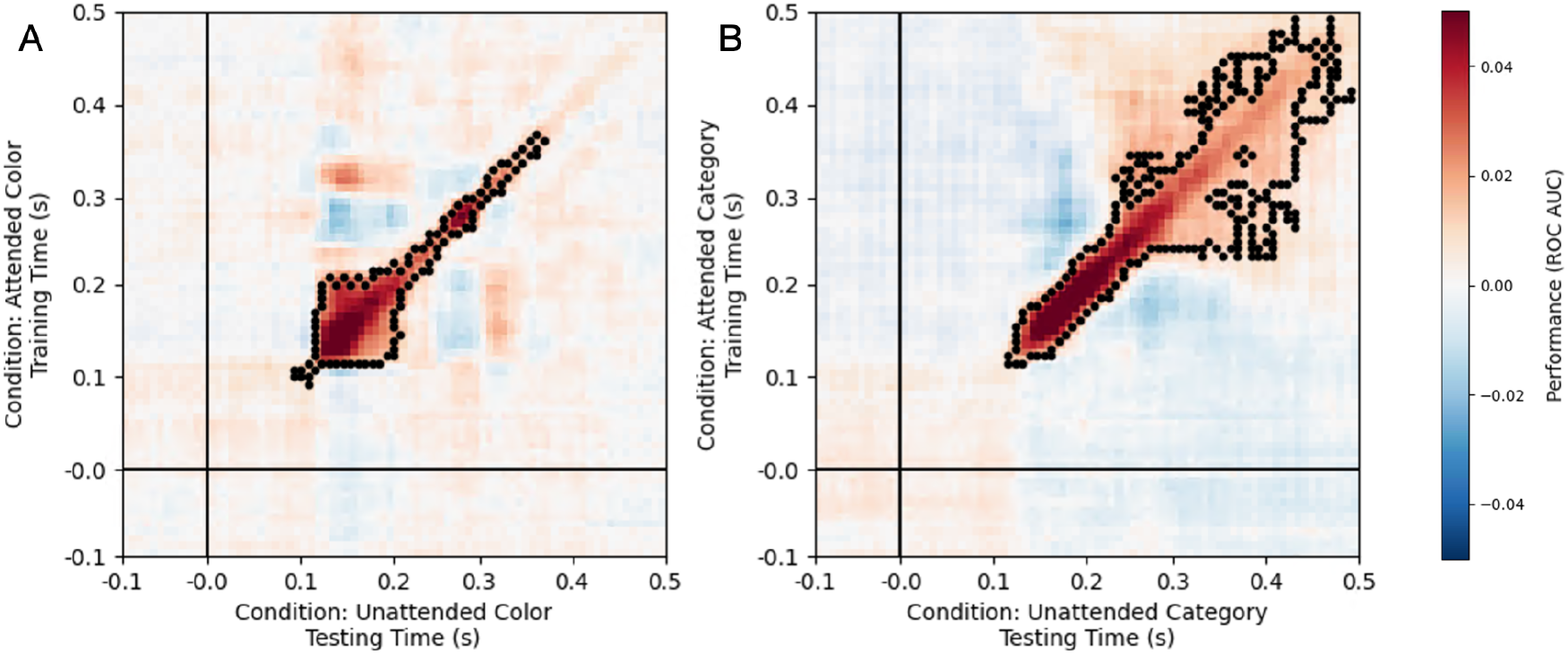
Decoding generalization between sessions averaged across participants. A) Decoding model of color identity. B) Decoding model of category identity. The x axis represents the time point of the testing set, recorded in the unattended session and the y axis represents the time point of the training set recorded in the attended session. Color reflects the AUC.

### Generalization across tasks

In order to determine whether representations are task-dependent we also used the model trained on the baseline block and tested it on the three independent tasks conducted in the same session. We detected significant clusters in both the color session (prediction task - range: 94-336 ms, p<0.001, perception task - range: 94-367 ms, p<0.001 and memory task - range: 102-360 ms, p<0.001; Figure 5A) and category session (prediction task - range: 109-500 ms, p<0.001, perception task - range: 109-445 ms, p<0.001 and memory task - range: 109-500 ms, p<0.001; Figure 5B).

**Figure 5.**
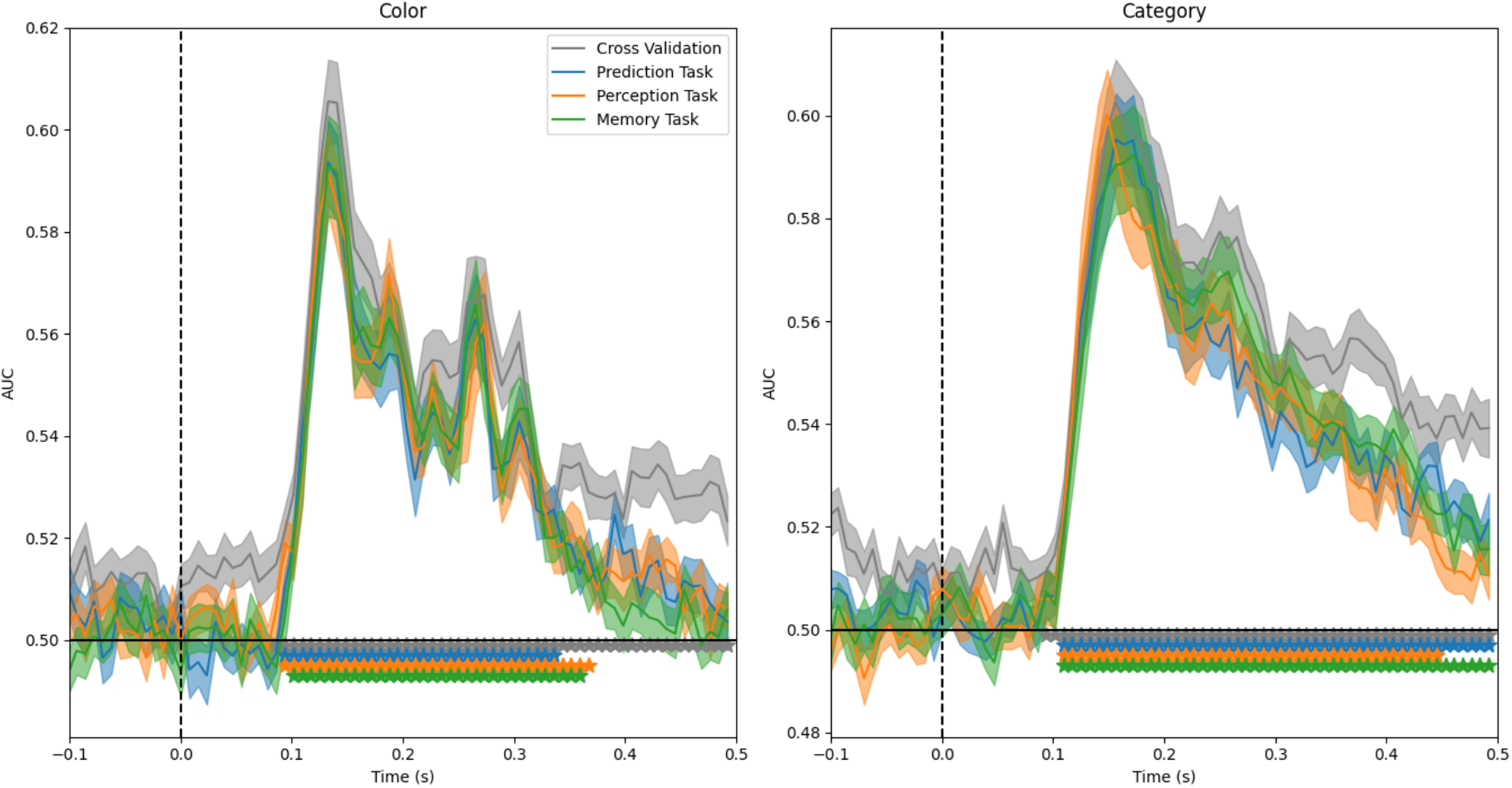
Decoding results of A) Color identity and B) Category identity. The x axis represents the time point relative to stimulus onset (represented by the dashed line). The Y axis represents AUC relatively to chance (marked by the black line and 0.5). The colored lines compared the results from the cross validation set to the other tasks (prediction, perception and memory tasks).

### Correlations between decoding accuracy and EEG signal

As demonstrated in the correlational figures (Figure 3), there was considerable variability among participants in decoding accuracy, which seem to be sustained across sessions, attentional conditions, and task. To understand whether properties of the ERP might covary with decoding accuracy we correlated it to the ERP signal (amplitude) and noise (trial variation). A marginally significant correlation between decoding accuracy and P1-N1 peak-to-peak amplitude, which did not cross the significance threshold after correction for multiple comparisons, was detected (Color: r = 0.39, p=0.033; Category: r= 0.39, p=0.031), see Figure 6A. A weak trend for a correlation between decoding accuracy and ERP trial-based variability was found for both category and color, which was not significant (Color: r = −0.24, p=0.201; Category: r= −0.33, p=0.078; Figure 6B).

**Figure 6.**
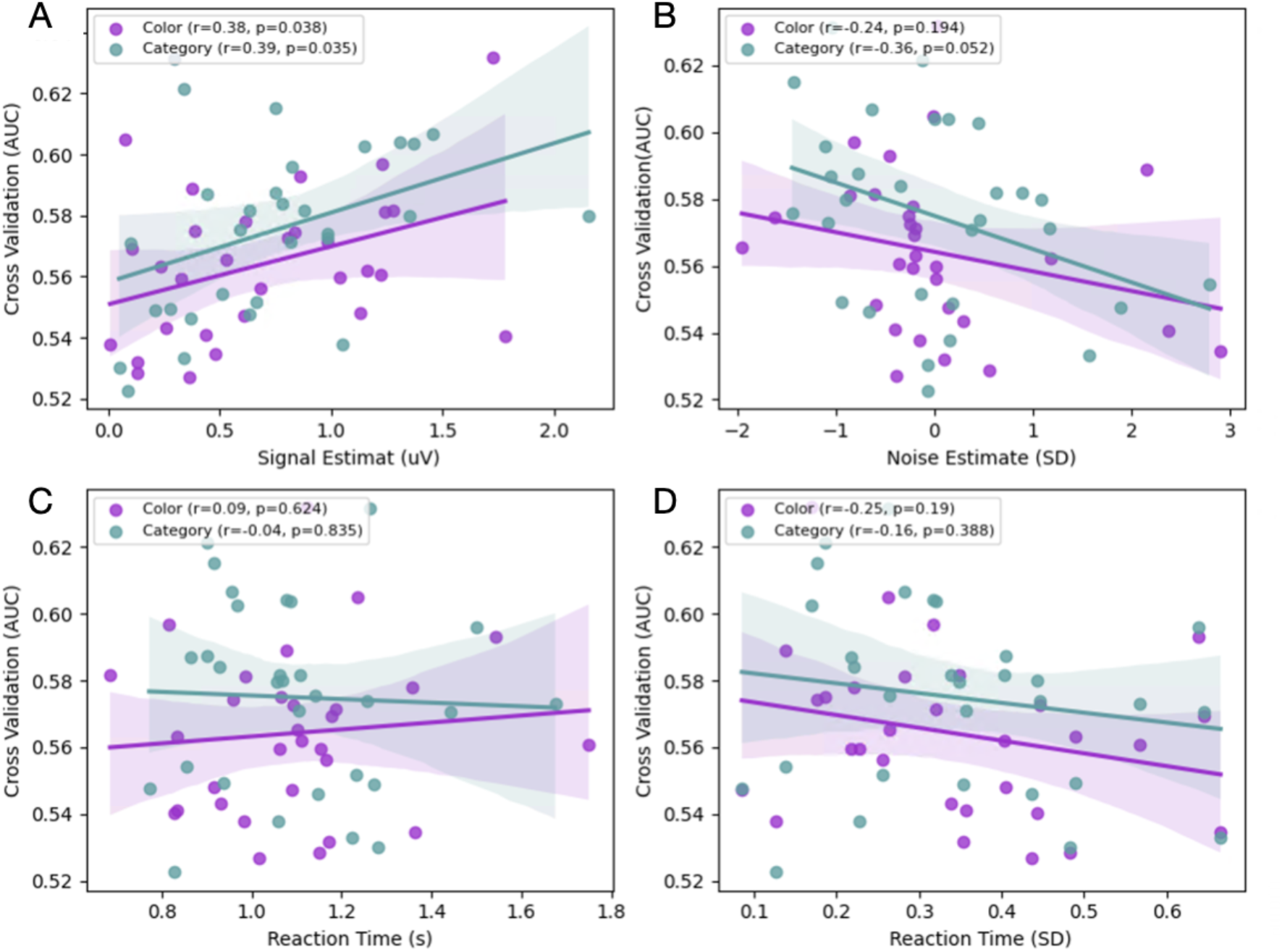
Scatter plot with linear trend of the cross-validation AUC at the perceptual time (0.1-0.3 seconds post stimulus onset, Y axis) separately for color and category (marked by color) per participant. A) with standardized EEG noise estimate (X axis). B) with peak-to-peak ERP amplitude (X axis) C) with behavioral reaction time in seconds (X axis). D) with the standard deviation of the reaction time in seconds (X axis)

### Correlations between decoding accuracy and task performance

In order to understand the relationship between task performance, as an estimate of task engagement and attention, and decoding accuracy we correlated the AUC with the mean and standard deviation of the reaction times in the baseline task (we did not test for correlation with behavior accuracy since it showed a very limited range due to a ceiling effect). No significant correlations were found (See Figure 6C-D), suggesting that behavioral performance in this design cannot account for the variability between participants.

## Discussion

Multivariate pattern analysis (MVPA) is increasingly used in neuroscience for decoding representations embedded in neuroimaging signals, providing a powerful tool for understanding the spatial and temporal distribution of different levels of information. However, to trust this measure, it is important to understand the reliability of decoding across sessions and conditions, and individual differences in decoding accuracy. In this study, we replicated robust above chance decoding of both color (Hajonides et. Al., 2021) and category (Contini et. Al., 2017) information from EEG signals, using a minimal preprocessing pipeline, without signal amplification (averaging to create “super-trials”, as in Iamshchinina et. Al., 2022) or dimensionality reduction. The temporal profiles of decoding mirrored known visual hierarchies: color information was decodable earlier and more briefly than category, consistent with early visual cortex engagement for color and a more distributed and long-lasting activation for category. We observed noticeable variability between participants in decoding accuracy. Individual decoding accuracies were found to be reliable across sessions, suggesting that decoding accuracy is not random but reflects trait-like characteristics. Furthermore, we show that decoding models of color and category can be generalized to other tasks and attentional conditions within the same session, as well as between sessions (1-7 days apart), pointing to stable underlying neural representations and robustness to variability in EEG measurement setup.

### People vary in a reliable way in the decodability of their brain signals

Our findings offer a closer examination of variability between participants in decoding results, which are rarely reported. While individual differences could be ascribed to or random fluctuations in cognitive states, or to measurement noise, especially considering the inherently low signal to noise ratios in the EEG (Goldenholz et. Al., 2009), the correlation analysis revealed that decoding accuracy is remarkably stable within participants. Decoding accuracy highly correlated between independent sessions separated by days, between attentional conditions (feature attended vs. not attended) and even between the decoding of color and categories. This result lends strong support for the reliability of decoding measures and suggests that ‘decodability’ is an individual trait.

This reliability raises the question of the sources of inter-subject variability. To try and elucidate this question, we examined the correlations between decoding accuracy on the one hand, and behavioral measurements, ERP signal amplitude, and ERP variability parameters on the other hand. We first hypothesized that participants who demonstrate a clearer signal (i.e. more prominent ERPs or lower inter-trial variability) might show higher decoding accuracy, as demonstrated in fMRI (Smith et. Al., 2011; Albers et. Al., 2018). The results of this analysis were inconclusive. We found a moderate, marginally significant positive correlation between the visual ERP amplitude and decoding accuracy. However, our measurements depended on a priori spatio-temporal regions of interest (the N1-P1 peak to peak at PO7/8), which may not an optimal measure of signal strength for every single subject, considering that the classifier is not limited in the same way. The analysis of inter-trial variability elicited only a trend for negative correlation with decoding accuracy, which seems to be associated with participants who demonstrated very high or very low trial-based variability. Future studies could investigate these issues with fewer decoding levels and higher number of subjects, allowing robust correlation analysis.

Since the classifier attempts to detect color and category identity a potentially important factor that could explain inter-subject variability is the accessibility of key neural regions to EEG sensors, such as V4 in the case of color decoding. Individuals vary in gyral and sulcal gross morphology and their relation to specific neuronal populations, which in turn project to scalp electrodes (Scrivener & Reader, 2022; Irimia et. Al., 2012; Ales et. Al., 2010). Thus, participants with optimal sensor coverage vis a vis individual functional neuroanatomy of visual areas involved in color and category processing might exhibit higher decoding performance. Testing this hypothesis will require a combination of fMRI with high density EEG and source.

Another source for individual differences could be motivation and engagement, or attentional state during the task. We hypothesized that participants with a higher RT or larger standard deviation of the RT, might have been less attentive and accordingly might show lower accuracy. However, we did not find a significant correlation between the reaction time or its standard deviation to decoding accuracy. This lowers the probability that the variability in decoding performance between participants is simply a byproduct of session-specific transient states like attention and motivation. Whereas some evidence for correlation between electrophysiological data decoding accuracy and trial specific behavioral performance has been reported (O’sullivan et. Al., 2015), as well as some studies linking accuracy (Williams et. Al., 2007) and RT (Grootswagers et. Al., 2018; Ritchie et. Al., 2015) to decoding accuracy of fMRI data, the current study does not support such a correlation. Our experimental design did not allow a trial-based behavioral estimate, as it prioritized the uninterrupted study of perceptual processes, with behavior only sampled sparsely. This limits our ability to examine whether trial by trial fluctuation of attention could affect the decodability. However, the lack of correlation between decoding accuracy and RT variability makes this a less plausible explanation, consistent with the results of (Desantis et. Al., 2020) using stimulus category level behavioral measures. Future studies may also investigate the possibility that participants with more vivid or distinctive visual representations may exhibit greater decoding accuracy, akin to individual differences in mental imagery ability (Pearson, 2019).

### Distinctive neural patterns for colors and categories are stable

The test-retest reliability of decoding accuracies does not entail, in and of its own, that the neural responses to different stimuli were similar across sessions or conditions. That is, if classifiers are trained de novo in every session, the decoding may be high in both sessions while relying on different discriminating patterns in each session. Our results show however that not only is the accuracy reliable, but so are the patterns of neural activity. Single participant decoding models of both color and category generalized well across sessions, tasks, and attentional conditions, suggesting that the underlying representations, extracted by the decoding model, were stable as well. This result provides strong validation to EEG decoding as a method for measuring neural representations of visual information. In the context of the current study, the results also highlight the specificity of the model to visual sensory representations, independent of the task. Taken together, the high between session accuracy correlation and pattern generalization support that decoding reflects stable participant-specific properties. This understanding might have important implications on experimental protocols and participants selection for decoding studies.

These findings also suggest that EEG-based decoding is not affected by representational drift in the timescales of days, which has been observed in single-unit recordings (Driscoll et al., 2022). The generalizability found in the current study aligns with recent work in fMRI showing that the pattern BOLD signals are stable over the time scale of days to weeks (Roth & Merriam, 2023). Taken together, the EEG and fMRI results suggest that representational drift is less evident at the neuronal population level measured by these methods. The cross-session generalization shown in the current study extends the fMRI results by showing not only spatial but also temporal stability. Future studies are required in order to estimate weather more significant drift may occur over longer time periods, as demonstrated by Roth & Merriam (2023).

### Implications for future MVPA studies

Our findings provide important implications for EEG studies employing decoding. Decoding analysis requires a large data set for training and meticulously balanced designs to avoid spurious results (Combrisson & Jerbi, 2015). A frequent limitation is the maximal session duration participants are able to endure and in which quality recording can be maintained. This leaves researchers with two options – collect more participants and measure different conditions on different groups (instead of within participant; e.g., Noah et. al, 2020 preform the control analysis on a different group of participants) or bring participants in for multiple sessions, to sample multiple conditions on the same group of participants. The current study suggests the first solution might be problematic, since when sampling two groups of participants, the control group could by chance include less decodable participants, potentially biasing the studies’ conclusions. We show that bringing in participants for multiple sessions, while maintaining a controlled setup, is a viable solution. This solution keeps the overall number of participants required to the study smaller while eliminating a significant risk of bias. Our findings correspond with a recent approach of “precision neuroscience” (Nebe et. Al., 2023). This approach suggests that acquiring more data from a smaller group of participants and using individualized analysis might reduce the variability between studies and increase replicability by improving the reliability of the measurements and statistical power. Precision neuroimaging has been applied to fMRI, sometimes using very small sample sizes and a participant-tailored analysis pipeline (Mueller et. Al. 2013, Laumann et al., 2015, Nettekoven et. al., 2024). Our findings, confirming the test-retest reliability and generalizability of decoding, suggest this approach might be applied to electrophysiology. By using two EEG sessions per participant, we could double the size of the training set and use independent training and test sets, while keeping all manipulations within participant and allowing for a within participant replication of the results. While measuring fewer subjects might hinder the generalizability of the results to the population, a balance may be stricken between the number of sessions and the number of participants. As suggested by Nebe et. al, this should be considered according to the specific study’s goals. While the results of the current study strongly supports the reliability of decoding of sensory content, the reliability in other domains like attention or mood should be investigated in future studies.

In summary, this study highlights the reliability and generalizability of MVPA applied to EEG data and the stability of individual differences in decoding performance. By demonstrating stable within-participant accuracy across sessions and tasks, we provide evidence that decoding performance reflects intrinsic neural properties rather than random variability or transient states. These findings contribute to a growing body of evidence supporting the robustness of MVPA in investigating neural representations across time and tasks, paving the way for broader applications in cognitive neuroscience. Future research should aim to disentangle the contributions of anatomical factors, representational strength, and cognitive traits to individual differences in decoding performance.

## Acknowledgments

We would like to thank the participants that participated in this study and the HCNL lab members of advice and support. C.F. is supported by the Mandel Foundation graduate fellowship. L.Y.D. is supported by the Jack H. Skirball research fund. This study was supported by US-Israel Binational Science Foundation grant 2024626. LYD is the founder and consultant of Innereye Ltd, a startup company in the field of neurotech. The company was not involved in this study.

## Footnotes

1 [CV decoding of color, category and their difference, difference between predicted and unpredicted CV decoding, cross-session color, cross-session category + 6 tasks]

